## Supplementary figures and images for "Pex30-dependent membrane contacts sites maintain ER lipid homeostasis"

### Fig S1

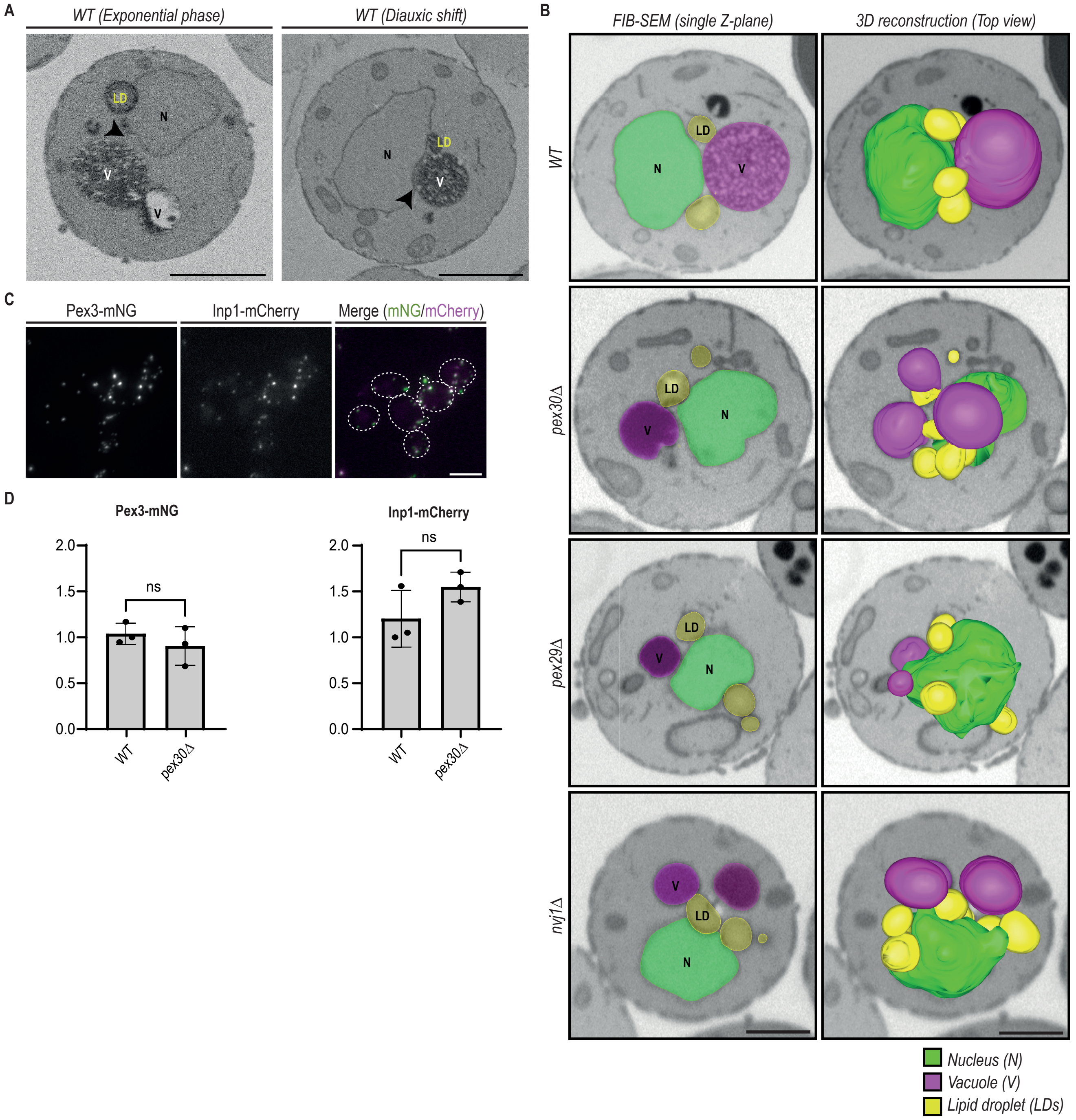

### Fig S2

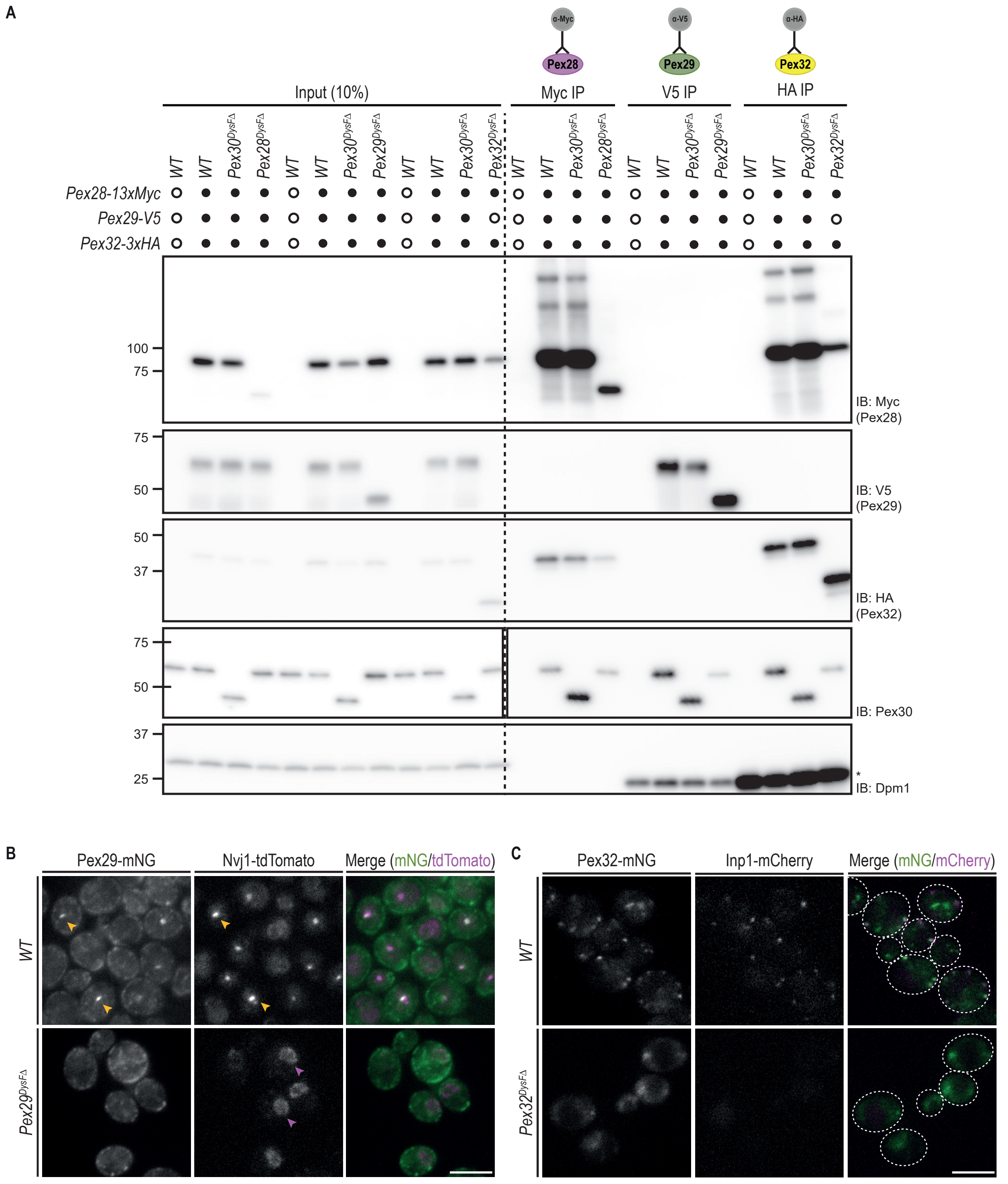

### Fig S3

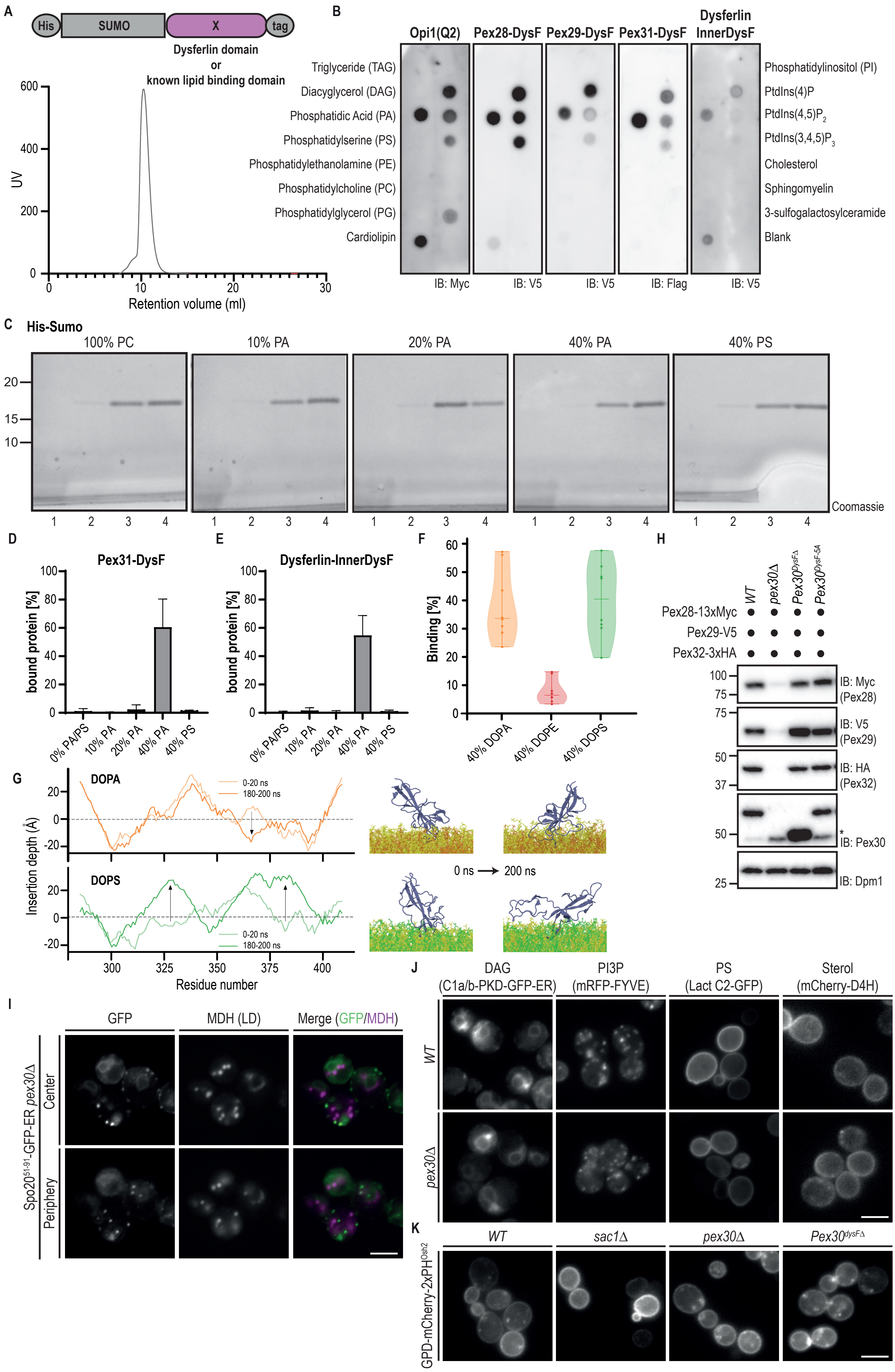

### Fig S4

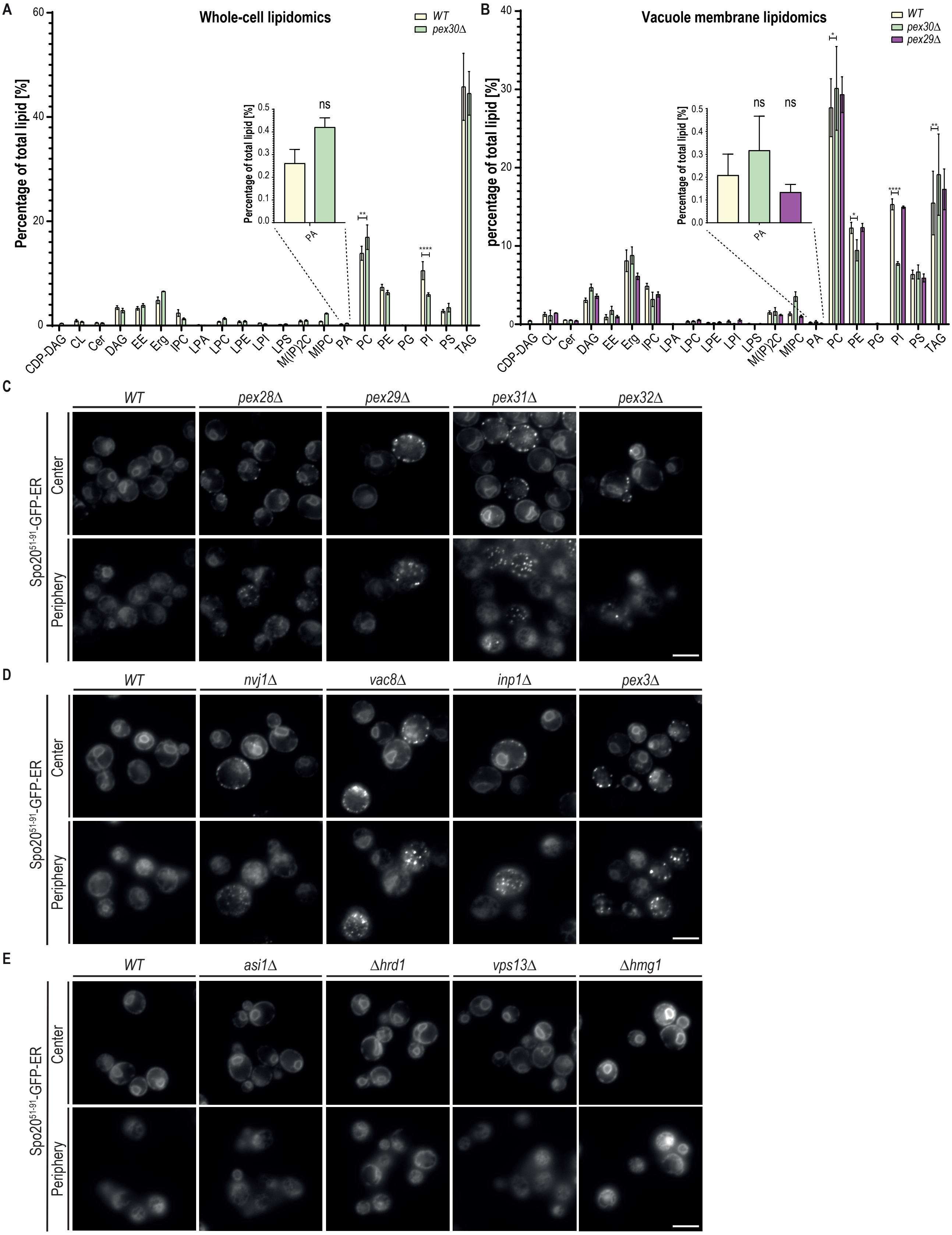

### Fig S5

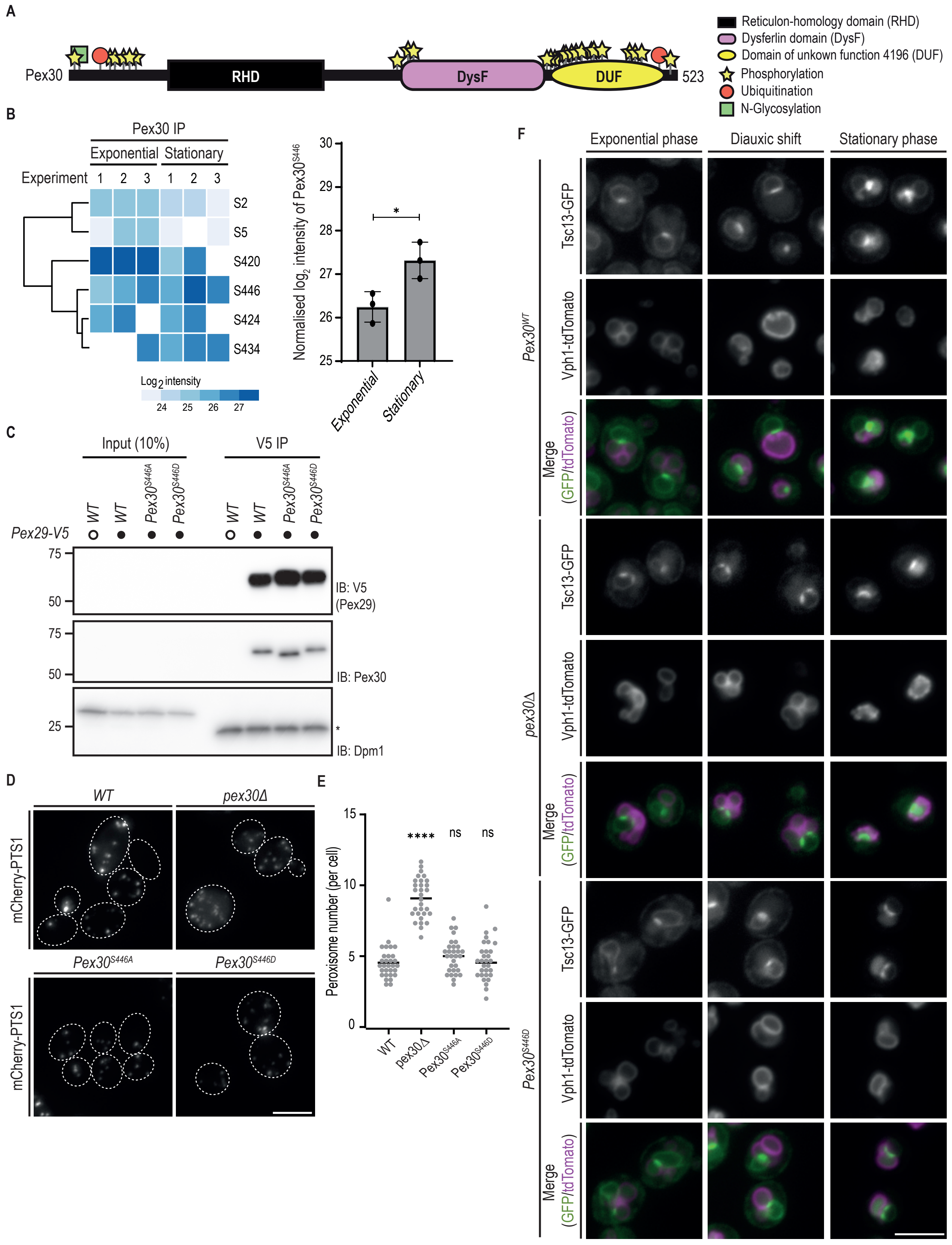
